# Beta hydroxybutyrate alters beta cell identity and function in human islets

**DOI:** 10.64898/2025.12.24.696385

**Authors:** Yun Suk Chae, Mandala Ajie, Twan J.J. de Winter, Esmay Hammink, Ezra van der Wel, Maaike Jantja Roodzant, Françoise Carlotti, Marten A. Engelse, Eelco J. P. de Koning

## Abstract

Fasting alters insulin secretion in humans and islet cell composition in mice. Fasting triggers ketogenesis, which increases beta hydroxybutyrate (BHB) levels. BHB is a signaling molecule that alters insulin secretion in primary islets. We hypothesized that BHB alters human islet cell composition and identity. Primary human islets were cultured with R-BHB or its non-metabolizable enantiomer, S-BHB. Human islets cultured with R-BHB, but not S-BHB, resulted in an increased C-peptide^-^glucagon^+^ to C-peptide^+^glucagon^-^ cell ratio, and an increased frequency of C-peptide^+^glucagon^+^ bihormonal cells and NKX6.1^+^glucagon^+^ cells. Single-cell transcriptomics revealed upregulation of alpha cell identity genes in R-BHB treated islet beta cells. Alterations in beta cell identity were accompanied by increased basal insulin secretion in response to low glucose, and a reduced insulin stimulation index in response to high glucose. The differential effects of the two BHB enantiomers on islet cell composition indicated that ketone metabolism is involved in islet cell identity change. Supporting this hypothesis, a subpopulation of beta cells in islets treated with R-BHB characterized by low ketolysis genes, *OXCT1* and *ACAT1*, did not show altered alpha and beta cell identity genes. Additionally, primary islets from type 2 diabetes donors, which exhibited reduced *OXCT1* and *ACAT1* expression relative to donors without diabetes, did not display altered islet cell composition and beta cell function from R-BHB treatment. Our findings uncover a novel role of BHB in modulating beta cell identity, with potential implications for islet function in states with chronically elevated ketone concentrations.

## Introduction

The rising prevalence of metabolic diseases, including obesity and diabetes, has stimulated growing interest in restrictive dietary strategies such as fasting [1, 2]. Fasting induces a range of catabolic processes, notably ketogenesis. Ketogenesis is a metabolic process in which the liver produces ketone bodies, beta hydroxybutyrate and acetoacetate, through fatty acid oxidation [3]. Beta hydroxybutyrate (BHB) can reach concentrations of 2 to 8 mmol/L during prolonged fasting [3], and serves as an alternative energy source for organs, such as the heart and kidney [4].

While short term fasting is associated with health benefits, such as weight loss and improved glycemic control [1, 2], studies on prolonged fasting in healthy adults show paradoxical effects of impaired glucose tolerance [5], reduced first-phase insulin secretion [6, 7], and increased insulin resistance [8, 9], prompting the term “pseudo-diabetes” to describe a fasted state [9]. Notably, fasting ketone concentrations are also positively associated with the incidence of type 2 diabetes [10, 11], while primary islets treated with BHB *in vitro* show increased basal insulin secretion [12–14], a hallmark feature of beta cells in type 2 diabetes [15–17].

Although the underlying mechanism behind BHB’s effects on beta cell function is unclear, fasting in mice induces marked changes in islet cell composition with a reduced proportion of beta cells and increased frequency of cells with mixed alpha and beta cell phenotypes within a few days [18]. Similarly, long-term ketogenic diet in mice resulted in reduced alpha and beta cell mass [19], and induced glucose intolerance [19, 20]. Whether elevated levels of BHB in ketogenic conditions have an influence on islet cell composition and identity is unknown.

Given the increasing interest in utilizing fasting and specifically ketones for metabolic regulation [21, 22], we conducted this study to understand the potential role of BHB in islet cell plasticity.

## Materials and Methods

### Human islets, pancreas, and cell line

Pancreases of deceased donors were obtained through the Eurotransplant multiorgan donation program. Research was only performed on pancreases that could not be used for clinical purposes and if research consent was available, according to Dutch national laws. Islet isolations took place at the Leiden University Medical Center (LUMC) using a semi-automated method as described earlier [23]. Donor characteristics are listed in the supporting information (SI appendix) table 1. All experiments were performed on islets from fractions that had ≥ 70% purity and were in culture for ≤ 3 days at the start of the experiment. Islets were cultured in CMRL 1066 medium (5.5 mmol/L glucose) containing 10% human serum, 20 μg/mL ciprofloxacin, 50 μg/mL gentamycin, 2 mmol/L L-glutamine, 10 mmol/L HEPES, and 1.2 mg/mL nicotinamide in 6 well-ultra-low attachment plates (Corning).

EndoC-BetaH1 (EndoC-BH1) cells were obtained from Univercell Biosolutions (Toulouse, France). Cells were seeded in ECM and fibronectin coated plates and cultured with low glucose DMEM medium (Invitrogen, USA) containing 10 mmol/L nicotinamide, 5.5 g/ml transferrin, 6.7 ng/ml selenite, penicillin-streptomycin, and 50 μmol/l beta-mercaptoethanol.

Powder of the R-enantiomer of beta hydroxybutyrate (R-BHB, Sigma Aldrich 54920), and the S-enantiomer, S-BHB (Sigma Aldrich 54925), were dissolved in aliquots of the islet and EndoC-BH1 medium. The pH of the BHB medium was adjusted using sodium hydroxide to the corresponding control medium. Islets were cultured in their assigned medium for 48-hours. EndoC-BH1 cells were cultured in their assigned medium for 48- and 96-hours. Medium change was performed every two days.

Pancreas tissue samples were fixed overnight in 4% formaldehyde, stored in 70% ethanol, and embedded in paraffin. Paraffin blocks were cut into 4-μm sections. Tissues from donors with a history of type 2 diabetes (n = 10) were compared with those without a history of diabetes (n = 10). Detailed information on donor backgrounds is outlined in SI appendix table 2.

### Immunofluorescence staining

Islets were fixed in 4% formaldehyde for 30 minutes at room temperature and washed with PBS. Islets were spun down in fluid agar, and the agar-embedded pellets were paraffinized. Paraffin blocks were cut in 4-μm sections. After deparaffinization, antigen retrieval was performed using 10 mmol/L sodium citrate buffer in a pressure cooker. Primary blocking step was performed using 5% goat serum for 1 hour. Sections were imaged using Zeiss LSM 900 microscopy. The investigators were blinded to the treatment conditions and donor backgrounds during all primary islet and pancreas imaging and quantifications. Further details on the antibodies used in this study and immunostaining quantifications can be found under the materials and methods section in the SI appendix.

### RNA isolation and quantitative PCR

Total RNA was obtained using RNeasy kit (Qiagen) according to the manufacturer’s protocol. cDNA was synthesized using M-MLV reverse transcriptase (Invitrogen) and quantitative PCR (qPCR) was performed using Light Cycler 480-II Real-time PCR system (Roche). Beta actin was used as the reference gene. The control group was set as 1 and statistical analysis was made between the fold change from R-BHB and S-BHB exposure. The primer sequences are listed in SI appendix table 3.

### Glucose-stimulated insulin secretion

Glucose-stimulated insulin secretion (GSIS) of the cultured islets was performed on the Perifusion System V5 (Biorep). Three hundred islet equivalents (IEQ) per treatment condition were preincubated for 1 hour in Krebs-Ringer bicarbonate buffer with 2 mmol/L glucose at 37°C. Islets were stimulated with 20 mmol/L glucose for 50 minutes, followed by a return to 2 mmol/L of glucose for 20 minutes, depolarized with KCl for 5 minutes, and finalized by 2 mmol/L of glucose for 10 minutes. Insulin concentrations in the perfusate were determined by ELISA (Mercodia). Average basal insulin secretion was calculated between t = 5 minutes till t = 15 minutes. The stimulation index for each measurement was calculated by setting average basal insulin secretion as the baseline. The average stimulation index and area under the stimulation index curve was calculated during 20 mM glucose exposure, which was between t = 20 minutes till t = 65 minutes. KCl depolarization was determined at t = 90 minutes. The secretory capacity of beta cells was considered intact if the stimulation index during KCl depolarization (KCl stimulation index) was ≥ 2.

### Baseline calcium flux imaging

Islets were incubated with 10 µM Cal-520 (Abcam, ab171868) and 0.04% Pluronic F-127 (Sigma-Aldrich, P2443) in Krebs–Ringer bicarbonate buffer (KRB) containing 2 mmol/L glucose for 90 minutes at 37°C, followed by 30 minutes at room temperature. Prior to imaging, the dye solution was replaced with fresh KRB containing 2 mmol/L glucose. Stained islets were transferred to a 96-wells plate previously coated with Matrigel (Corning) diluted 1:22 in KRB buffer. The islets were allowed to attach to the Matrigel for 30 minutes at 37°C. For coating, diluted Matrigel was added to the wells and incubated for 1h at 37°C, after which the solution was aspirated to leave a thin layer. Imaging was performed using an EVOS M7000 (Thermo Fisher Scientific), with images acquired every 5 seconds for a total duration of 3 minutes.

Image analysis was performed using ImageJ. Regions of interest (ROIs) were created for each individual islet, and the fluorescence intensity was measured over time. Baseline stability for each islet was quantified as the absolute value of the slope of fluorescence intensity over time, with values closer to 0 indicating more stable baseline calcium flux.

### Single-cell transcriptomics on R-BHB treated primary islets

Islets isolated from 3 donors were treated with or without R-BHB for 48-hours and dissociated into single-cells using a 0.025% trypsin (Gibco) and 10 mg/mL DNase (Pulmozyme, Genentech, San Francisco, CA, USA) mix. Dispersed islet cells were filtered through a 20 µm cell strainer (Pluriselect, 43-50020-03) to remove clumped cells. The viability and cell counts of the single-cells were quantified using NucleoCounter NC-200 (Chemometec). The single-cell samples had a viability ranging between 81% and 97%. Cells were frozen in Cryostor (Sigma, C2874) and stored in liquid nitrogen prior to 10x genomics sequencing. Single-cell transcriptomics was performed by Single-cell Discoveries (The Netherlands) according to standard 10X Genomics 3’gene expression protocols. Sequencing results were mapped using Cell Ranger count.

Data analysis involving R-BHB treated islets was performed in R version 4.3.3 using Seurat package version 5.3.0 [24]. To remove ambient RNA, SoupX (version 1.6.2) was applied with an ambient RNA threshold fixed at 25%. Doublets were removed using DoubletFinder (version 2.0.4). Cells with total UMI between 500-50,000 reads, gene counts > 1,000 reads, and mitochondrial gene reads < 25% were included in the analysis. Cells were annotated using known islet gene markers as previously described [25]. Differential gene expression (DGE) was done using the FindMarkers function from Seurat that is based on Wilcoxon Rank Sum test. DGE analysis included genes that were expressed in at least 25% of the cells from any group. Genes were considered significantly altered if they had an Benjamini-Hochberg-corrected *p* value of < 0.05. Gene set enrichment analysis was performed using clusterProfiler package (version 4.10.1) with Molecular Signature Database (MSigDB) Hallmark gene set collection.

Composite beta and alpha cellular identity gene scores were calculated using Ucell package (version 2.6.2). UCell creates composite scores based on Mann-Whitney U statistics using user-provided marker gene signatures [26]. Alpha and beta identity scores for endocrine cells were calculated using identity marker genes identified by van Gurp et al [27].

Python package Scanpy (version 1.9.3) and the CollecTRI model within the package decoupleR (version 2.0.6) were used to perform single-cell transcription factor activity (TFA) analysis. First, an activity inference with a univariate linear model based on the expressed genes in the dataset was run. After fitting, the t-values of the slopes were used as the transcription factor activity scores. Differential TFA was performed using the rankby_group in decoupleR with the t-test_overestim_var option. Transcription factors with a Benjamini-Hochberg adjusted p-value below 0.05 were considered differentially active.

### Type 2 diabetes single-cell transcriptomics dataset

12 published SMART-seq and SMART-seq2 datasets of human pancreatic islets were integrated. FASTQ files were downloaded from Wang et al. (GSE83139) [28], Xin et al., (GSE81608) [29], Enge et al. (GSE81547) [30], Segerstolpe et al. (E-MTAB-5061) [31], Lawlor et al. (GSE86469) [32], Li et al. (GSE73727) [33], Camunas-Soler et al. (GSE124742) [34], Dai et al. (GSE164875) [35], Liu et al (GSE270484) [36], Marquez-Curtis et al. (Human Cell Atlas) [37], and SMART-seq and Patch-seq datasets from the Human Pancreas Analysis Program (HPAP) [38]. Further details can be found under the materials and methods section in the SI appendix.

### Statistical analysis

Data are expressed as mean ± SD or ± SEM, as stated in the figures. Statistical significance of differences between two groups were determined by a *t* test. For multiple groups, statistical significance was first assessed using one-way ANOVA, and then Fisher’s least significant difference test was performed. *p* < 0.05 was considered statistically significant. Statistical analysis was performed on GraphPad Prism 10.2.3.

## Results

### R-BHB alters primary human islet cell composition

We performed immunostaining for C-peptide and glucagon on human islets cultured with or without R-BHB for 48-hours (Fig. 1A). R-BHB treated islets showed reduced frequency of C-peptide^+^glucagon^-^ cells (71.3±6.4% vs 55.8±5.6%, CTRL vs R-BHB, *p* = 0.0009; Fig. 1B), and increased frequency of C-peptide^-^glucagon^+^ cells (25.7±6.6% vs 39.5±5.5%, CTRL vs R-BHB, *p* = 0.0017; Fig. 1C) and C-peptide^+^glucagon^+^ cells (3.0±1.7% vs 4.7±2.3%, CTRL vs R-BHB, p = 0.012; Fig. 1D, 1E). R-BHB treated islets showed a 95% increase in the C-peptide^-^glucagon^+^ to C-peptide^+^glucagon^-^ cell ratio (0.37±0.12 vs 0.72±0.16, CTRL vs R-BHB, *p* = 0.0015; Fig. 1F).

**Figure 1:**
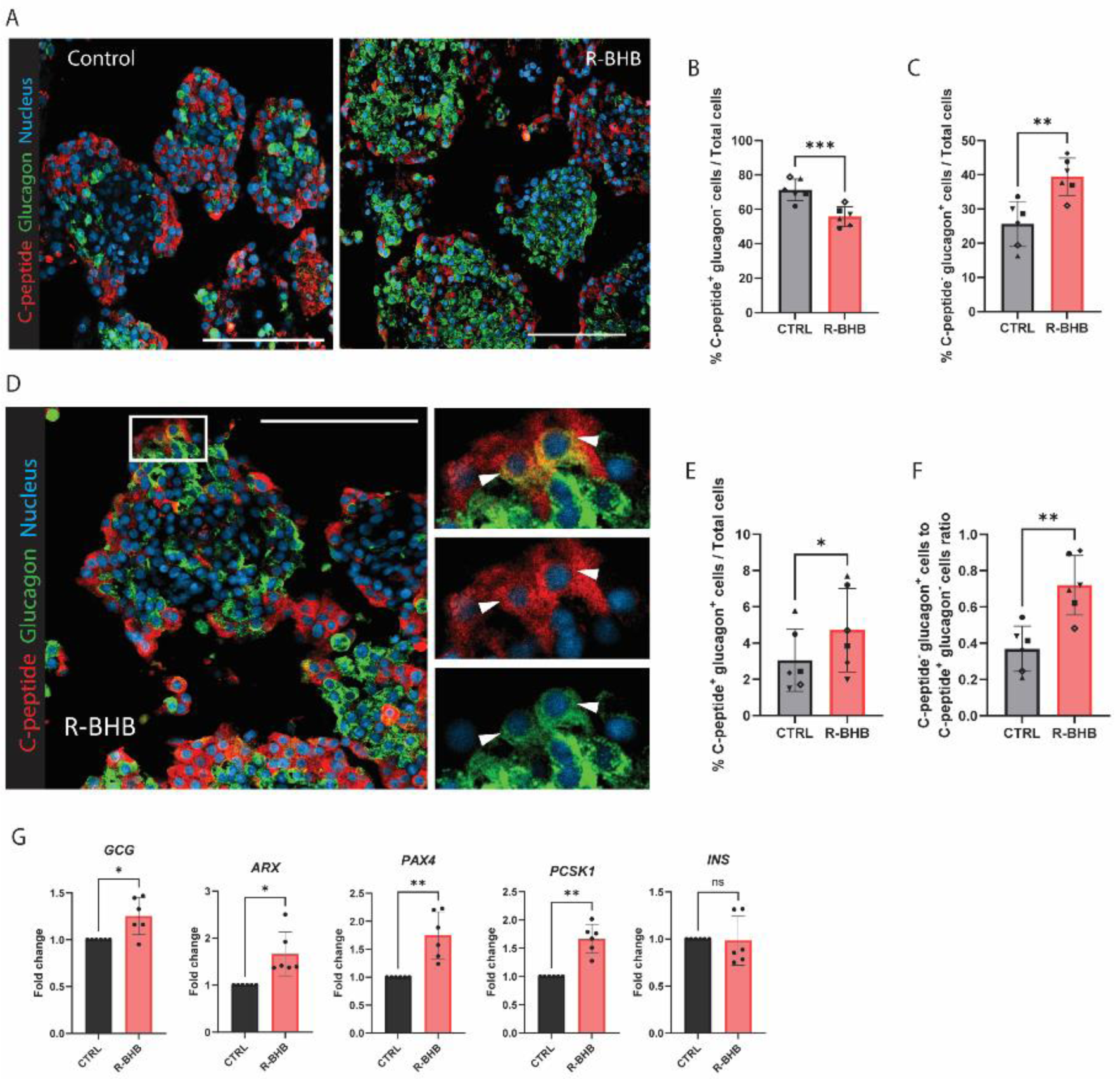
R-BHB alters primary human islet cell composition. *A*: Representative immunostainings for C-peptide (red) and glucagon (green). Image on the left are islets cultured in control medium, and on the right are islets cultured in R-BHB medium. *B, C*: Quantification of C-peptide^+^glucagon^-^ and C-peptide^-^glucagon^+^ cells. *D*: Representative immunostaining for C-peptide^+^glucagon^+^ bihormonal cells in islets cultured in R-BHB medium. In the insert, white arrows indicate the nuclei of C-peptide^+^glucagon^+^ bihormonal cells. *E*: Quantification of C-peptide^+^glucagon^+^ bihormonal cells. *F:* Quantification of the C-peptide^-^glucagon^+^ and C-peptide^+^glucagon^-^ cells ratio. *G*: qPCR analysis of islets cultured with or without R-BHB. n = 6 donors. Data are represented as mean ± SD. Paired Student’s t-test (*B*, *C*, *E*-*G*) was used to assess statistical significance. (\**p* < 0.05, \*\**p* < 0.01, \*\*\**p*<0.001). ns = non-significant. Scale bar 100 µm.

Quantitative PCR (qPCR) analysis showed upregulation of alpha cell markers *GCG* (1.25±0.20 fold change, *p* = 0.027) and *ARX* (1.66±0.46 fold change, *p* = 0.032), and beta cell markers *PAX4* (1.75±0.42 fold change*, p* = 0.0071) and *PCSK1* (1.67±0.25 fold change, *p* = 0.0065) in R-BHB treated islets (Fig. 1G). There was no difference in *INS* expression levels following R-BHB treatment (Fig. 1G).

To assess whether the effects of BHB treatment are R-BHB specific, an enantiomer of BHB that cannot be directly metabolized, S-BHB, was included as an additional control. Islets treated with S-BHB for 48-hours did not show alterations in their cell composition (SI appendix Fig. 1A, 1B, 1C, 1D).

We then examined whether R-BHB is cytotoxic to islets or induces proliferation in order to explain changes in islet cell composition. Islets cultured in 0.5, 5, and 10 mM R-BHB showed comparable viability to those in the control medium (SI appendix Fig. 1E, 1F). There was no difference in beta cell apoptosis and overall cell apoptosis from 5 mM R-BHB treatment (SI appendix Fig. 1G, 1H, 1I). We found no differences in alpha cell proliferation or overall tissue proliferation between the two groups (SI appendix Fig. 1J, 1K, 1L).

### Single-cell transcriptomics identifies increased alpha cell identity gene expression in R-BHB treated islet beta cells

To investigate the endocrine cell populations that underwent identity changes, we performed single-cell transcriptomic analysis on three donor islet preparations treated with or without R-BHB for 48-hours (Figure 2A). After quality controls and filtering, a total of high-quality 40,344 non-treated and 37,550 BHB treated human pancreatic cells were obtained. Unsupervised clustering of the pancreatic cells showed 10 populations (Figure 2B). The endocrine cell populations, which included alpha, beta, delta, gamma, and polyhormonal cells were distinguished using established identity markers [27] (Fig. 2C).

**Figure 2:**
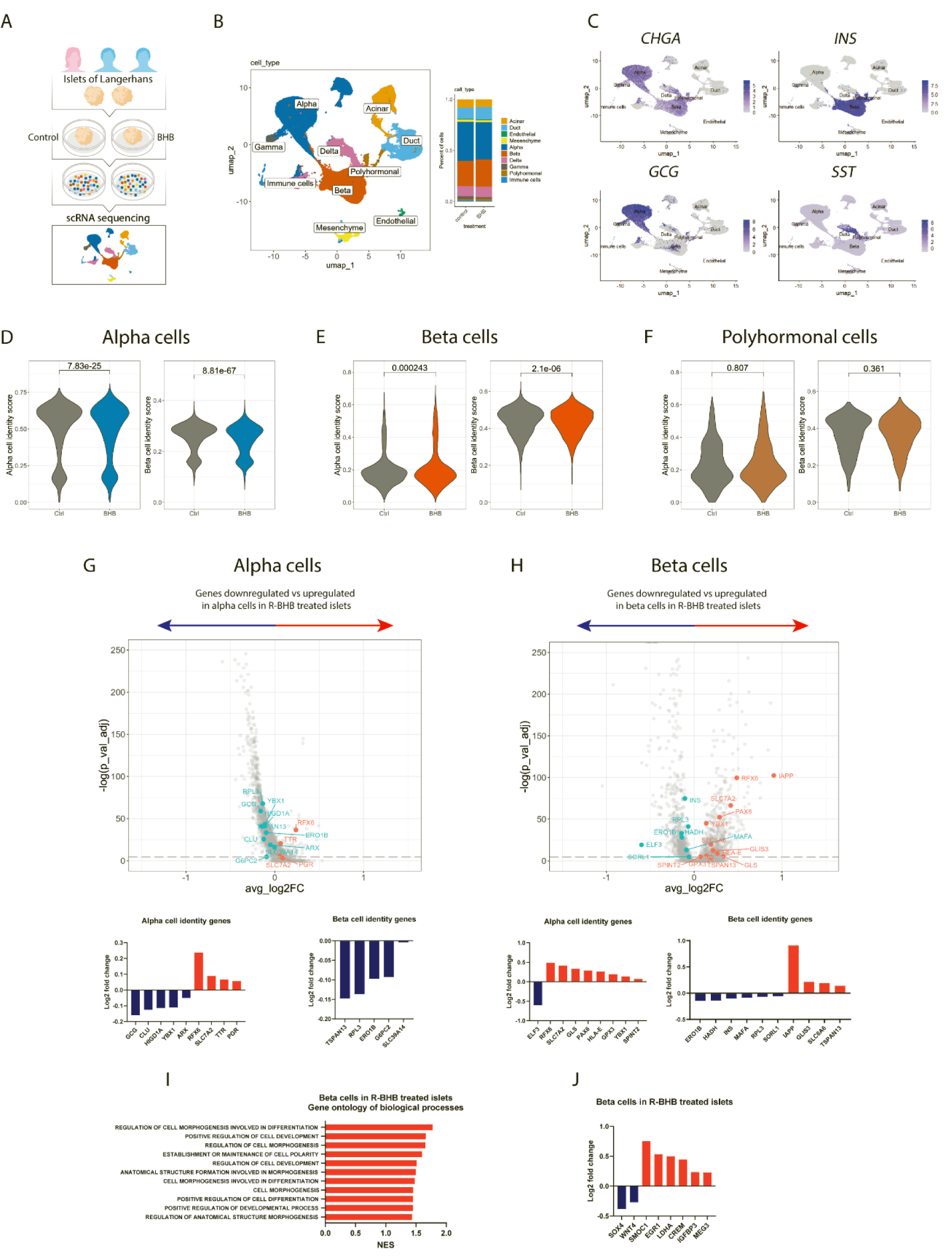
Single-cell transcriptomics identifies increased alpha cell identity gene expression in R-BHB treated beta cells. *A*: Experimental scheme for single-cell transcriptomics of R-BHB treated primary human islets from 3 donor islet preparations. *B*: UMAP and cell composition of annotated pancreatic cell types. *C*: UMAP of the distribution of key endocrine identity markers. *D, E, F*: Comparisons of alpha and beta cell identity scores in alpha, beta, and polyhormonal cells in islets treated with and without R-BHB. *G*: Differential gene expression analysis of alpha and beta cell identity genes in alpha cells in R-BHB treated islets. *H*: Differential gene expression analysis of alpha and beta cell identity genes in beta cells in R-BHB treated islets. *I*: Gene ontology enrichment analysis of beta cells in R-BHB treated islets. *J*: Differentially expressed genes involved in beta cell dedifferentiation, cell development and identity, stress response and mass maintenance, and disallowed gene in beta cells in R-BHB treated islets. Numbers on the violin graphs represent adjusted *p*-values.

We first examined overall changes in the alpha and beta cell identity scores in alpha, beta, and polyhormonal cell populations from islets treated with R-BHB. R-BHB treated islets contained alpha cells with lower alpha and beta cell identity scores (Fig. 2D) and beta cells with higher alpha cell and lower beta cell identity scores compared to control (Fig. 2E). Polyhormonal cells did not show alterations in alpha and beta cell identity scores (Fig. 2F).

We next performed a differential gene expression (DGE) analysis of alpha and beta cells in R-BHB treated islets to identify alterations in the top 10 identity markers, 10 transcription factors, and 10 cell surface protein-encoding genes for alpha and beta cells, as reported by van Gurp et al [27]. Alpha cells in R-BHB treated islets showed alpha cell marker downregulation of *GCG, CLU, HIGD1A*, *YBX1*, and *ARX* and upregulation of *RFX6*, *SLC7A2*, *TTR*, and *PGR,* and beta cell marker downregulation of *TSPAN13, RPL3, ERO1B, G6PC2,* and *SLC39A14* (Fig. 2G). Beta cells in R-BHB treated islets showed beta cell marker downregulation of *ERO1B, HADH, INS, MAFA, RPL3, SORL1*, and upregulation of *IAPP, GLIS3, SLC6A6,* and *TSPAN13* (Fig. 2H), and alpha cell marker upregulation of *RFX6, SLC7A2, GLS, PAX6, HLA-E, GPX3, YBX1,* and *SPINT2* (Fig. 2H). Gene set enrichment (GSE) analysis on R-BHB treated islets revealed upregulation of gene sets in beta cells associated with cell fate and morphogenic regulation (Fig. 2I). Additional DGE analysis of beta cells in R-BHB treated islets showed upregulation of beta cell dedifferentiation gene *SMOC1*, alterations in genes associated with beta cell development and identity maintenance (*SOX4*, *WNT4*, and *MEG3*), upregulation of genes associated with stress responses and beta cell mass maintenance (*EGR1, CREM,* and *IGFBP3*), and upregulation of the disallowed gene, *LDHA* (Fig. 2J).

### Sub-clustering analysis identifies a sub-population of beta cells expressing *GCG,* and R-BHB increases the frequency of glucagon expressing beta cells

As R-BHB treated islets exhibited upregulation of multiple alpha cell markers in beta cells, we hypothesized that beta cells may be the primary contributors to the observed changes in islet cell composition following R-BHB treatment. Therefore, we performed an additional unsupervised sub-clustering analysis on the beta cell population, which revealed three subpopulations: Beta 1, Beta 2, and Beta 3 (Fig. 3A). All three subpopulations expressed *INS,* while Beta 3 also expressed *GCG* (Fig. 3B). Further characterization of the beta cell subpopulations in terms of their alpha and beta cell markers showed that Beta 1 and Beta 2 displayed high levels of key beta cell identity genes, such as *MAFA* in Beta 1 and *NKX6.1* in Beta 2, while Beta 3 was characterized by high levels of key alpha cell markers, such as *ARX* and *IRX2* (Fig. 3C).

**Figure 3:**
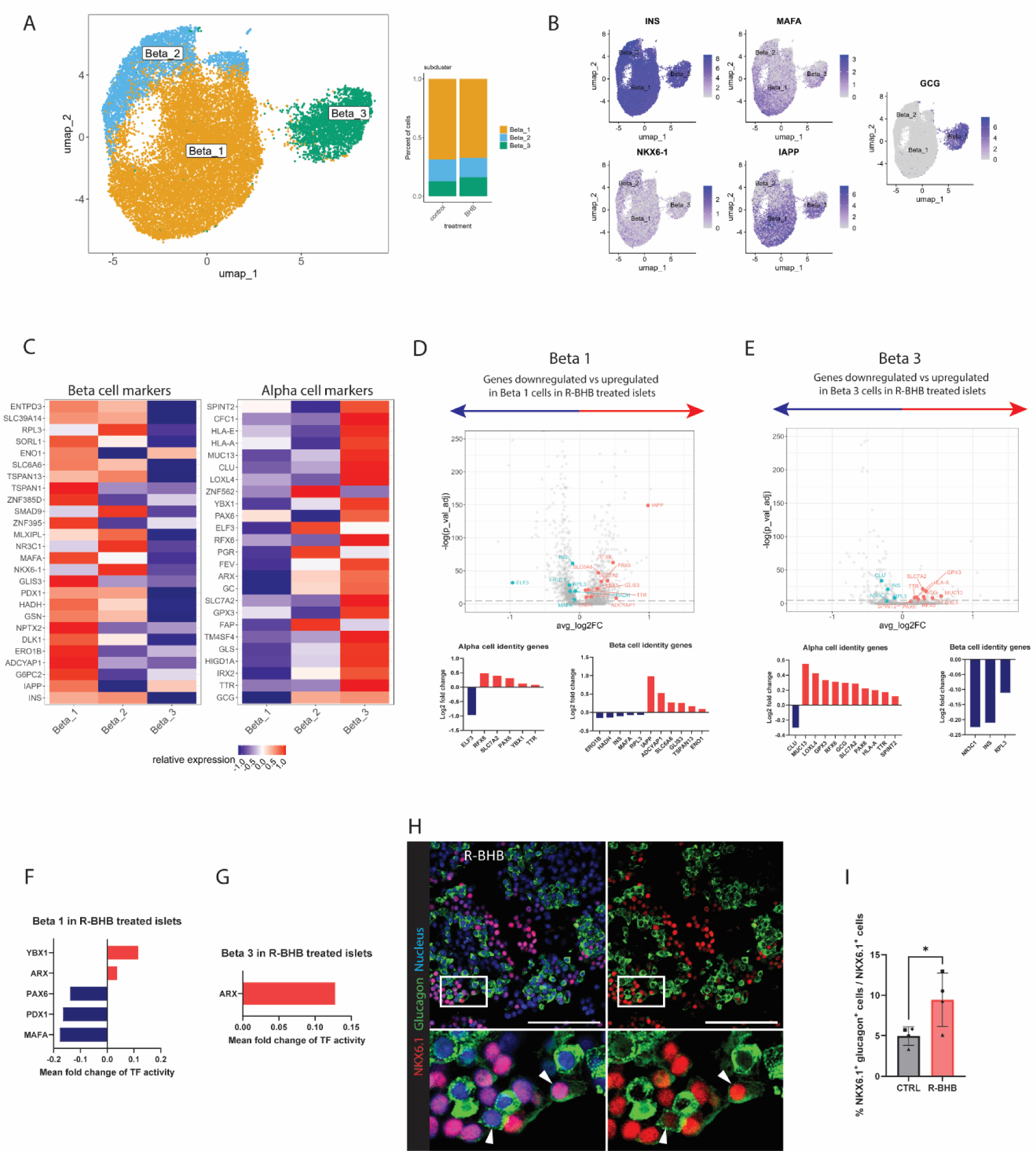
Sub-clustering analysis identifies a sub-population of beta cells expressing *GCG*, and R-BHB increases the frequency of glucagon expressing beta cells. *A*: UMAP of the proportion of beta cell subpopulations. B: UMAP of the distribution of key beta cell identity genes and *GCG* in the beta cell subpopulations. *C*: Heatmap of the relative expression of beta and alpha cell markers among the 3 beta cell sub-populations. *D, E*: Differential gene expression analysis of alpha and beta cell identity genes in Beta 1 and Beta 3 sub-populations in R-BHB treated islets. *F, G*: Transcription factor activity analysis of Beta 1 and Beta 3 subpopulations in R-BHB treated islets. *H*: Representative immunostaining for glucagon (green), NKX6.1 (red), and nucleus (blue) of islets cultured in R-BHB medium. In the insert, the white arrows indicate the nuclei of NKX6.1^+^glucagon^+^ cells. *I*: Quantification of NKX6.1^+^glucagon^+^ cells in total NKX6.1^+^ cells. n = 4 donors. Paired Student’s t-test (*I*) was used to assess statistical significance. (\**p* < 0.05). Scale bar 100 µm.

We next examined the DGE of alpha and beta cell identity markers in beta cell subpopulations from R-BHB treated islets. Beta 1 showed downregulation of beta cell markers *ERO1B, HDAH, INS, MAFA,* and *RPL3* and upregulation of alpha cell markers *RFX6, SLC7A2, PAX6, YBX1,* and *TTR* (Fig. 3D). Beta 2 showed no alterations in the expression levels of both alpha and beta cell identity genes. Beta 3 showed downregulation of beta cell markers *NR3C1, INS,* and *RPL3,* while alpha cell markers *MUC13, LOXL4, GPX3, RFX6, GCG, SLC7A2, PAX6, HLA-A, TTR,* and *SPINT2* were upregulated (Fig. 3E).

To better understand the transcriptional regulators underlying the changes in beta cell identity following R-BHB treatment, we examined the transcription factor activity, which we infer based on the expression of their target genes. Beta 1 in R-BHB treated islets showed decreased activity of *PAX6*, *PDX1,* and *MAFA*, and increased activities of *YBX1* and *ARX* (Fig. 3F). Beta 3 in R-BHB treated islets showed increased activity of *ARX* (Fig. 3G).

To identify beta cells with alpha cell identity following R-BHB treatment, we selected NKX6.1 as a beta cell specific marker [39, 40], as neither qPCR nor single-cell transcriptomics analyses showed alterations in *NKX6.1* expression in R-BHB treated islets, islet beta cells, and islet alpha cells (SI appendix Fig 2A). Immunostaining revealed an 88% increase in NKX6.1^+^glucagon^+^ cells following R-BHB treatment (5.0±1.1% vs 9.4±3.3%, CTRL vs R-BHB, *p* = 0.034; Fig. 3I, 3J). Additionally, EndoC-BetaH1 (EndoC-BH1) cells, an immortalized human beta cell line, treated with R-BHB showed upregulation of *GCG* (1.23±0.17 fold change, *p* = 0.0447; SI appendix Fig. 2B) after 96-hours, but not in S-BHB treated cells (SI appendix Fig 2B).

### R-BHB treated islets display increased basal insulin secretion and basal calcium flux, and reduced insulin stimulation index

To assess whether alterations in beta cell identity translates to functional changes, we performed a glucose stimulated insulin secretion (GSIS) test (Fig. 4A). R-BHB treated islets showed an increased average basal insulin secretion (*p* = 0.0411; Fig. 4B). Corrected for basal insulin secretion (Fig. 4C), R-BHB treated islets showed a reduced average stimulation index during 20 mM glucose exposure (*p* = 0.0153; Fig. 4D) and reduced area under the stimulation index curve (*p* = 0.0256; Fig. 4E). Islets treated with or without R-BHB showed similar beta cell secretory capacity to KCl (KCl stimulation index, CTRL vs R-BHB 12.7±8.3 vs 8.24±4.53, *p* = 0.08). R-BHB treated islets showed increased calcium baseline slope, indicating greater baseline calcium fluctuations (*p* = 0.0307; Fig. 4F, 4G).

**Figure 4:**
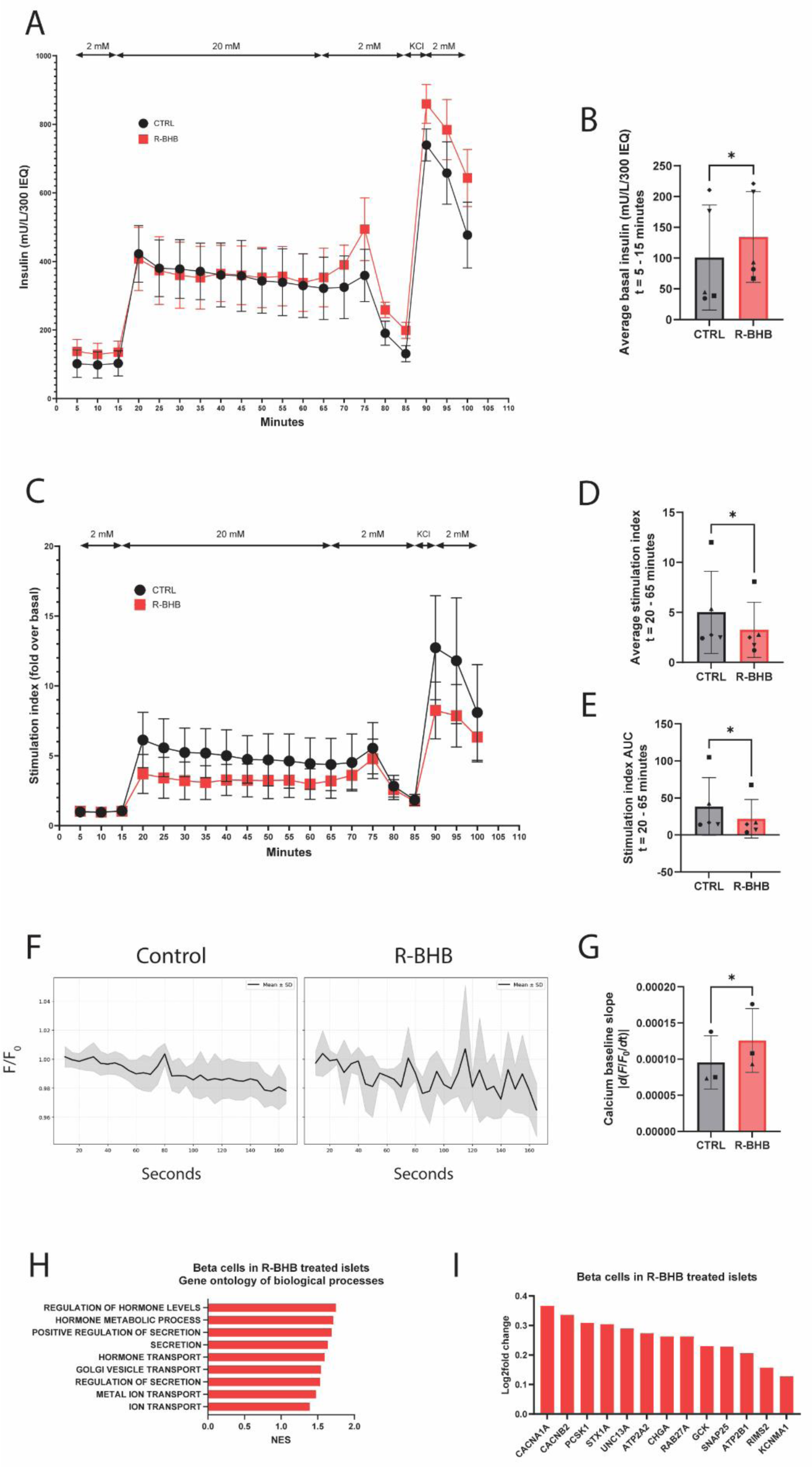
R-BHB treated islets display increased basal insulin secretion and basal calcium flux, and reduced insulin stimulation index. *A*: Glucose stimulated insulin secretion of 300 islet equivalents (IEQ) treated with or without R-BHB. *B*: Comparison of the average basal insulin secretion (2 mM glucose, between t = 5 and t = 15 minutes). *C*: The stimulation index profiles of islets treated with or without R-BHB. The stimulation index was calculated by setting the average basal insulin secretion (between t = 5 and t = 15 minutes) as 1. *D*: Comparison of the average stimulation index during 20 mM glucose exposure (between t = 20 and t = 65 minutes). *E*: Comparison of the insulin stimulation index areas under the curve (AUC) (between t = 20 and t = 65 minutes). n = 5 donors. *F*: Representative calcium fluorescent intensity measurements over 3 minutes from 2 mM glucose exposure. Gray areas represent mean ± SD. *G*: Comparison of the average calcium baseline slope of islets exposed to 2 mM of glucose for 3 minutes. n = 3 donors. *H*: Gene ontology enrichment analysis of pathways associated with hormone secretion and regulation and ion transport in R-BHB treated beta cells. *I*: Differentially expressed genes involved in calcium signaling, membrane excitability, granule formation, insulin processing, and glucose sensing in R-BHB treated beta cells. Data are represented as mean ± SEM (*A*, *C*) and as mean ± SD (*B, D*-*G*). Ratio t-test (*B*, *D*, *E*, *G*) was used to assess statistical significance. (\**p* < 0.05).

GSE analysis in beta cells of R-BHB treated islets revealed upregulation of pathways associated with hormone secretion and regulation and ion transport (Fig. 4H), while DGE analysis in beta cells showed upregulation of genes associated with increased beta cell calcium signaling (*CACNA1A, CACNB2, ATP2B1, RIMS2, KCNMA1*), membrane excitability (*ATP2A2*), granule formation (*STX1A, UNC13A, CHGA*), insulin processing (*PCSK1, RAB27A, SNAP25*), and glucose sensing (*GCK*) (Fig. 4I).

### Human islets have the capacity to metabolize ketones and alterations in beta cell identity from R-BHB treatment are associated with ketolytic enzymes

As S-BHB, the non-metabolizable enantiomer, did not induce islet cell composition change nor upregulate *GCG* expression in EndoC-BH1 cells, we hypothesized that metabolism of BHB is involved in beta cell identity change. Therefore, we examined whether human islets have the capacity to metabolize ketones by identifying the following key facilitators of ketone metabolism: monocarboxylic acid transporter 1 and 2 (MCT1, MCT2), 3-hydroxybutyrate dehydrogenase 1 (BDH1), 3-oxoacid CoA-transferase 1 (OXCT1), and acetyl-CoA acetyltransferase (ACAT1) (Fig. 5A). Immunostainings showed MCT1, BDH1, and ACAT1 expression in alpha cells and beta cells (Fig. 5B, 5D, 5F). MCT2 expression was low in islet cells (Fig. 5C). OXCT1 expression was high in beta cells and low in alpha cells (Fig. 5E).

**Figure 5.**
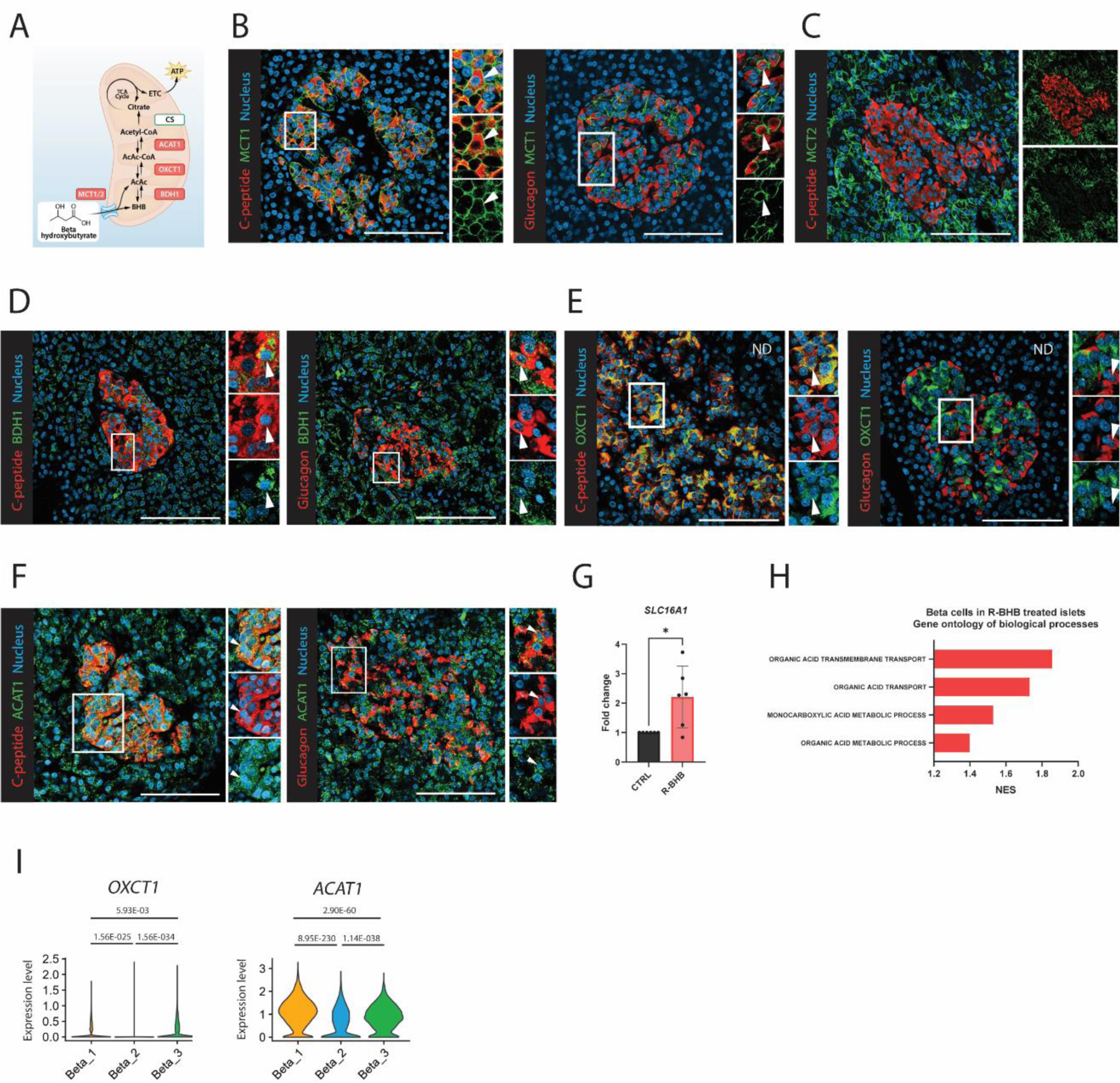
Human islets have the capacity to metabolize ketones and alterations in beta cell identity from R-BHB treatment are associated with ketolytic enzymes. *A*: Graphic representation of the ketone metabolism pathway. *B*: Representative immunostainings for MCT1 (green), C-peptide (red) or glucagon (red), and nucleus (blue) in donor pancreas. In the insert, white arrows indicate an example of C-peptide^+^MCT1^+^ cell and glucagon^+^MCT1^+^ cell. *C*: Representative immunostaining for MCT2 (green), C-peptide (red), and nucleus (blue) in donor pancreas. The insert shows low expression of MCT2 in beta cells and the area within the islet. *D*: Representative immunostainings for BDH1 (green), C-peptide (red) or glucagon (red) and nucleus (blue) in donor pancreas. In the insert, white arrows indicate the nuclei of C-peptide^+^BDH1^+^ cell and glucagon^+^BDH1^+^ cell. *E*: Representative immunostainings for OXCT1 (green), C-peptide (red) or glucagon (red) and nucleus (blue) in donor pancreas. In the insert, white arrows indicate the nuclei of C-peptide^+^OXCT1^+^ cell and glucagon^+^OXCT1^-^ cell. *F*: Representative immunostainings for ACAT1 (green), C-peptide (red) or glucagon (red), and nucleus (blue) in donor pancreas. In the insert, white arrows indicate the nuclei of C-peptide^+^ACAT1 ^+^ cell and glucagon^+^ACAT1^+^ cell. *G*: qPCR analysis of *SLC16A1* expression in R-BHB treated islets. n = 6 donors. *H*: Gene ontology enrichment analysis of R-BHB treated beta cells shows upregulation of pathways associated with BHB metabolism. *I*: Violin plot of *OXCT1* and *ACAT1* expression in the beta cell subpopulations. Numbers on the violin graphs represent adjusted *p*-values. Data are represented as mean ± SD. Paired Student’s t-test (*G*) was used to assess statistical significance. (\**p* < 0.05).

We then examined changes in genes involved in BHB metabolism following R-BHB treatment. R-BHB treated islets showed upregulation of *SLC16A1,* which encodes MCT1 (2.21±1.05 fold change, *p* = 0.0372; Fig. 5G). R-BHB treated islets showed enrichment for organic acid uptake and monocarboxylic acid metabolism pathways in beta cells (Fig. 5H). Further characterization of the beta cell subpopulations revealed that Beta 2, the subpopulation that did not show altered alpha or beta cell identity genes from R-BHB treatment, exhibited lower ketolysis genes *OXCT1* and *ACAT1* [3] compared to both Beta 1 and Beta 3 subpopulations (Fig. 5I).

### Beta cells from individuals with type 2 diabetes display lower capacity for ketone metabolism, and R-BHB treatment does not alter the cell composition of type 2 diabetes islets

Given the prominent protein expression of OXCT1 in beta cells, we investigated whether beta cell OXCT1 protein expression is altered in beta cells of type 2 diabetes (T2DM) donors compared to those from donors without a history of diabetes (ND). Beta cells from T2DM pancreas donors showed reduced expression of OXCT1 (% C-peptide^+^ OXCT1^+^/C-peptide^+^, 65±25% vs 37±24%, ND vs T2DM, *p =* 0.02; Fig. 6A, 6B).

**Figure 6.**
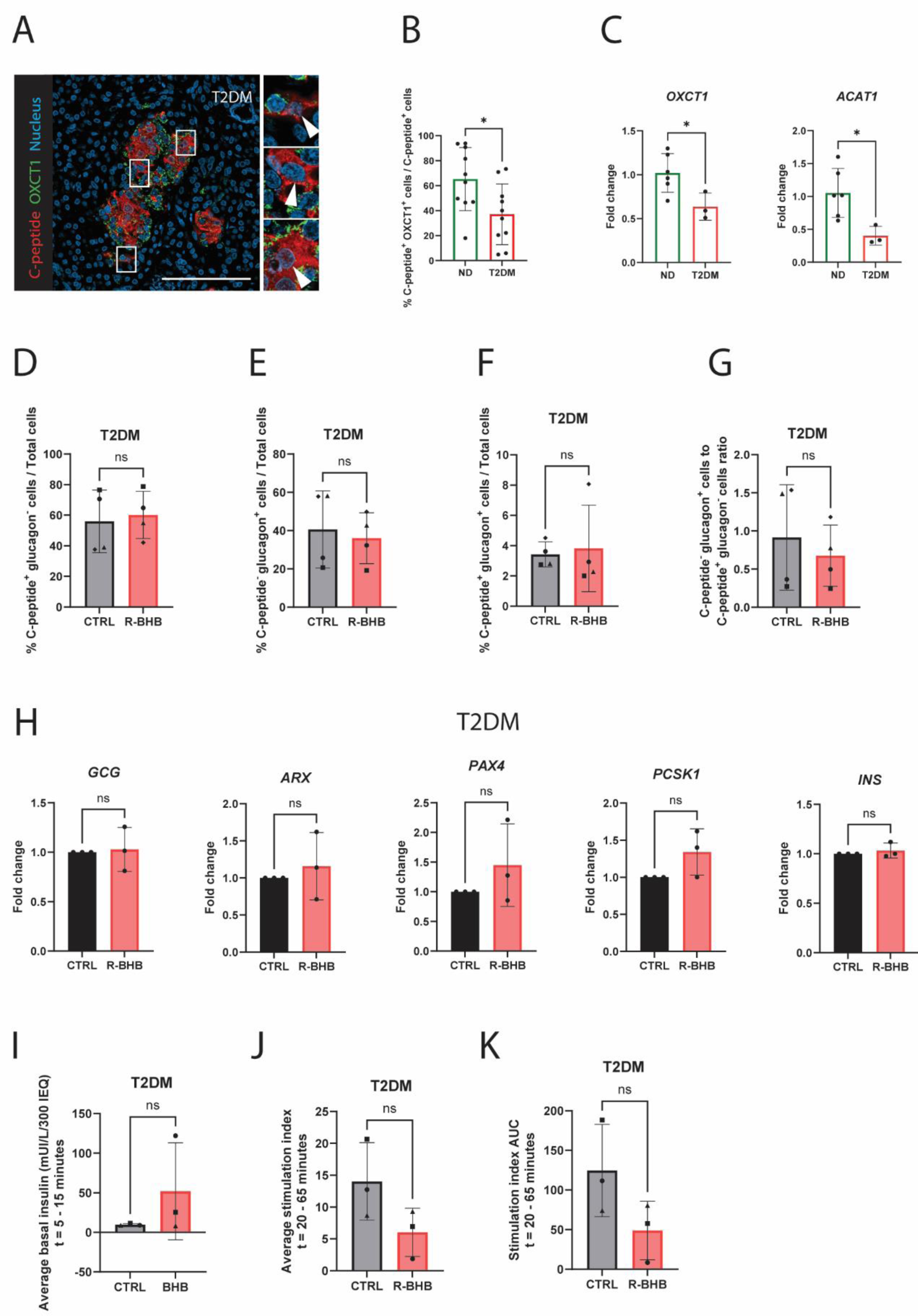
Islets from individuals with type 2 diabetes display lower capacity for ketone metabolism and do not alter their cell composition from R-BHB treatment. *A*: Representative immunostainings for OXCT1 (green), C-peptide (red), and nucleus (blue) in donor pancreas with a history of type 2 diabetes (T2DM). In the insert, white arrows indicate the nuclei of C-peptide^+^OXCT1^-^ cells *B*: Quantification and comparison of C-peptide^+^OXCT1^+^ cells in C-peptide^+^ cells in pancreatic islets from donors without diabetes (ND) and T2DM donors. n = 10 for ND pancreas donors and n = 10 for T2DM pancreas donors. *C*: qPCR analysis and comparison of *OXCT1* and *ACAT1* expression in islets from ND (n = 6) and T2DM (n = 3) donors. *D, E, F, G*: Quantification of C-peptide^+^glucagon^-^ and C-peptide^-^glucagon^+^ cells, C-peptide^+^glucagon^+^ bihormonal cells, and C-peptide^-^glucagon^+^ and C-peptide^+^glucagon^-^ cells ratio from donor islets with T2DM treated with or without R-BHB. n = 4 donors. *H*: qPCR analysis of T2DM islets cultured with or without R-BHB. *I*: Comparison of the average basal insulin secretion (2 mM glucose, between t = 5 and t = 15 minutes) from a GSIS of 300 IEQ from T2DM donors treated with or without R-BHB. *J*: Comparison of the average stimulation index (20 mM glucose, between t = 20 and t = 65 minutes). *K*: Comparison of the insulin stimulation index areas under the curve (AUC) (between t = 20 and t = 65 minutes). n = 3 donors. Data are represented as mean ± SD. Unpaired Student’s t-test (*B*, *C*), paired Student’s t-test (*D-G*), and ratio t-test (*I*-*K*) were used to assess statistical significance. (\**p* < 0.05). ns = non-significant. Scale bar 100 µm.

We also examined the expression of genes involved in monocarboxylic acid and ketone metabolism in T2DM beta cells by using publicly available single-cell transcriptomics datasets comprising of 90 ND donors and 35 T2DM donors. DGE analysis revealed a total of 65 genes involved in the monocarboxylic acid and ketone body metabolism were altered in T2DM beta cells, including the downregulation of *BDH1*, a key ketone metabolism enzyme involved in the interconversion of BHB and acetoacetate [3] (SI appendix Fig. 3).

From these observations, we hypothesized that primary T2DM islets have reduced plasticity in response to BHB. Primary islets from donors with HbA1c levels above the diagnostic threshold for diabetes (≥ 6.5% (48 mmol/mol Hb)) showed reduced expression of *OXCT1* (*p* = 0.0319; Fig. 6C) and *ACAT1* (*p* = 0.0199; Fig. 6C) compared to ND donors. T2DM islets treated with R-BHB did not show alterations in their islet cell composition (Fig. 6D, 6E, 6F, 6G), *GCG, ARX, PAX4, PCSK1,* and *INS* expression (Fig. 6H), average basal insulin secretion (Fig. 6I), average insulin stimulation index during 20 mM glucose exposure (Fig. 6J), and stimulation index area under the curve (Fig. 6K). T2DM islets treated with or without R-BHB showed similar beta cell secretory capacity to KCl (KCl stimulation index, CTRL vs R-BHB, 22.6±10.7 vs 8.9±5.6, *p* = 0.2).

## Discussion

Our study demonstrates that the ketone body, beta hydroxybutyrate (BHB) alters primary human islet cell composition and promotes beta cell transition towards an alpha-cell-like state. Ketolytic enzymes in beta cells are associated with these changes.

Clinical studies on fasting report that insulin and glucose levels decline, whereas glucagon and BHB levels rise [41–43]. In mice, prolonged fasting reduces beta cell proportions [18]. BHB treated primary human islets showed increased frequency of C-peptide^-^glucagon^+^ cells relative to C-peptide^+^glucagon^-^ cells, seemingly mirroring the physiological shift in hormone demands and reduced beta cell proportions found during fasting. These alterations in islet cell composition seem to be driven by a shift in beta cell identity towards an alpha-cell-like state, as evidenced by increased frequency of C-peptide^+^glucagon^+^ cells, NKX6.1^+^glucagon^+^ cells, and upregulation of alpha cell markers in BHB treated islet beta cells. Islet cells with a mixed alpha and beta cell identity are often described as transitional cells that rise in response to stress conditions [44–46]. BHB, therefore, may act as an adaptive signal to fasting conditions, prompting beta cells to adopt alpha-cell-like features.

The increased C-peptide^-^glucagon^+^ cells to C-peptide^+^glucagon^-^ cells ratio, increased frequency of cells with mixed alpha and beta cell identity, increased basal insulin secretion, and reduced stimulation index from BHB treatment are also features reminiscent of islets from people with a type 2 diabetes history [15, 16, 44, 47]. Elevated circulating ketones are associated with impaired insulin secretion, worsening of hyperglycemia, and increased incidence of type 2 diabetes [10, 11]. Furthermore, prolonged fasting induces insulin resistance and reduces insulin secretion [7, 9], while long-term ketogenic diet in mice results in glucose intolerance [19, 20]. An additional link between BHB and diabetes is observed in a distinct subtype of diabetes known as ketosis prone diabetes, which is characterized by the onset of diabetes with ketoacidosis without an identifiable precipitating cause [48, 49]. Individuals with ketosis prone diabetes exhibit impaired utilization of BHB for energy and elevated fasting BHB levels [48], suggesting that their pancreatic islets may be chronically exposed to elevated levels of BHB. Lastly, our findings may be relevant for understanding the development of malnutrition-related diabetes (Type 5 diabetes) that is associated with impaired insulin secretion [50]. The persistent nutritional ketosis due to chronic undernutrition [51] may act as a metabolic stressor that alters pancreatic beta cell function and identity. Whether there is a causal relationship between BHB and islet dysfunction associated with diabetes is unknown. However, the phenotypical similarities between islets exposed to BHB and in type 2 diabetes raise the question whether a mild chronic ketogenic state has adverse effects on beta cell health, and highlight the need for further *in vivo* studies.

BHB is a chiral molecule that exists as two enantiomers: R-BHB and S-BHB. Of these, only R-BHB is endogenously synthesized and directly metabolized [3]. The differential effects of the two BHB enantiomers on islet cell composition and beta cell identity led us to hypothesize that ketone metabolism is involved in the maintenance of beta cell identity. The central steps of ketone metabolism involve the uptake of BHB via the monocarboxylic transporter 1 (MCT1), interconversion of beta hydroxybutyrate and acetoacetate by beta hydroxybutyrate dehydrogenase 1 (BDH1), conversion of acetoacetate to acetoacetyl-CoA via the rate-limiting enzyme 3-oxoacid CoA-transferase 1 (OXCT1), and finally interconversion of acetoacetyl-CoA to acetyl-CoA by acetyl-CoA acetyltransferase 1 (ACAT1) [3]. Here we show protein expression of MCT1, BDH1, OXCT1, and ACAT1 in beta cells indicating the presence of key components necessary for BHB utilization. Of these, MCT1, encoded by *SLC16A1*, has traditionally been considered a disallowed gene in beta cells [52, 53]. However, a recent study has reported MCT1 protein expression in some human islet beta cells [54], corroborating our own findings. Moreover, we observed that *SLC16A1* expression was upregulated in R-BHB treated islets, suggesting that ketone availability may regulate its expression.

In support of a potential role of ketolytic enzymes in changes in islet cell identity, a subpopulation of beta cells (Beta 2) in islets treated with R-BHB, characterized by low expression of ketolysis genes *OXCT1* and *ACAT1*, did not show altered expression of alpha and beta cell identity genes. This finding was strengthened by observations in islets from donors with type 2 diabetes (T2DM). In T2DM islets, the proportion of C-peptide^+^ cells expressing OXCT1 was reduced, consistent with a previous report of lower OXCT1 protein abundance in T2DM islets [55]. Additionally, primary T2DM islets expressed lower *OXCT1* and *ACAT1*, while T2DM beta cells showed reduced expression of multiple genes involved in monocarboxylic acid and ketone metabolism, most notably *BDH1*. Collectively, these observations indicated a diminished capacity for BHB utilization in T2DM islets and led us to hypothesize that T2DM islets may be less able to alter their identity in response to BHB. Supporting this hypothesis, primary islets from donors with HbA1c levels above the diagnostic criteria for T2DM (6.5%) treated with R-BHB did not show changes in their islet cell composition, islet cell identity genes, and beta cell function.

The underlying mechanisms driving reduced expression of ketolytic enzymes in beta cells from T2DM donor islets are unclear. One possibility is that this reflects a broader feature of mitochondrial dysfunction in T2DM beta cells [56, 57]. Alternatively, it may represent an adaptive response aimed at preserving beta cell identity by limiting beta cell transition towards an alpha-cell-like state and help maintain residual beta cell functionality under ketogenic conditions.

A limitation of this study is that we do not show direct evidence that ketone metabolism alters islet cell composition and identity. It is also not clear whether altered islet cell identity can be translated to a (patho)physiological state of increased ketones, but our data reveal a previously unrecognized effect of ketones on human islet cell identity.

In conclusion, alterations in human islet cell composition and beta cell identity by beta hydroxybutyrate may have short- and long-term consequences for beta cell function and health.

## Data availability

Data available on reasonable request from the authors.

## Acknowledgements

The authors thank the donors and their families for providing valuable research material. The authors also thank the islet isolation team, Dirk-Jan Cornelissen, Corine Vermeulen-Hangelbroek, Evelien Rossenberg, Maaike Hanegraaf, and Bas Brinkhof for performing the islet isolations and providing research material.

## Funding

EJPdK has received funding from the DON Foundation, the Dutch Diabetes Research Foundation, JDRF, the Bontius Foundation, RegMedXB, and the Novo Nordisk Foundation Center for Stem Cell Medicine reNEW (NNF21CC0073729).

## Notes

### Competing Interest Statement

The Leiden University Medical Center has filed a patent application on a finding in this work.

