## Supporting information appendix for "Beta hydroxybutyrate alters beta cell identity and function in human islets"

**SI appendix table 1: Donor backgrounds for primary human islets**

| **Islet preparation** | **1** | **2** | **3** | **4** | **5** | **6** | **7** | **8** |
| --- | --- | --- | --- | --- | --- | --- | --- | --- |
| Donor age (years) | 30 | 57 | 75 | 46 | 73 | 60 | 72 | 69 |
| Donor sex (M/F) | M | M | M | M | M | F | F | M |
| Donor BMI (kg/m^2^) | 34 | 30 | N.D. | 25 | 33 | 24 | 27 | 29 |
| Donor HbA_1c_ | 32 mmol/mol (5.1%) | 38 mmol/mol (5.6%) | N.D. | 31 mmol/mol (5%) | N.D. | 39 mmol/mol (5.7%) | 34 mmol/mol (5.3%) | 33 mmol/mol (5.2%) |
| Origin/source of islets | LUMC | LUMC | LUMC | LUMC | LUMC | LUMC | LUMC | LUMC |
| Islet isolation centre | LUMC | LUMC | LUMC | LUMC | LUMC | LUMC | LUMC | LUMC |
| Donor history of diabetes | No | No | No | No | No | No | No | No |
| Donor cause of death | Brain death | Circulatory death | Brain death | Circulatory death | Circulatory death | Brain death | Circulatory death | Circulatory death |
| Warm ischaemia time | 0 minutes | 19 minutes | 0 minutes | 15 minutes | 20 minutes | 0 | 21 minutes | 24 minutes |
| Cold ischaemia time | 4 hours 52 minutes | 14 hours | N.D. | 5 hours 48 minutes | 16 hours 23 minutes | 16 hours 57 minutes | 13 hours 20 minutes | 13 hours 1 minute |
| Estimated purity (%) | 95 | 80 | 95 | 95 | 90 | 88 | 95 | 98 |
| Total culture time | 5 days | 5 days | 3 days | 3 days | 3 days | 3 days | 3 days | 3 days |
| Additional notes | N.D. | N.D. | N.D. | N.D. | N.D. | N.D. | N.D. | N.D. |

| **Islet preparation** | **9** | **10** | **11** | **12** | **13** | **14** | **15** | **16** |
| --- | --- | --- | --- | --- | --- | --- | --- | --- |
| Donor age (years) | 70 | 47 | 54 | 67 | 51 | 63 | 73 | 70 |
| Donor sex (M/F) | M | M | M | M | F | F | F | M |
| Donor BMI (kg/m^2^) | 27 | 27 | 24 | 26 | 20 | 16 | 26 | 25 |
| Donor HbA_1c_ | 38 mmol/mol (5.6%) | 36 mmol/mol (5.4%) | N.D. | 36 mmol/mol (5.4%) | 40 mmol/mol (5.8%) | 39 mmol/mol (5.7%) | 34 mmol/mol (5.3%) | 37 mmol/mol (5.5%) |
| Origin/source of islets | LUMC | LUMC | LUMC | LUMC | LUMC | LUMC | LUMC | LUMC |
| Islet isolation centre | LUMC | LUMC | LUMC | LUMC | LUMC | LUMC | LUMC | LUMC |
| Donor history of diabetes | No | No | No | No | No | No | No | No |
| Donor cause of death | Circulatory death | Circulatory death | Circulatory death | Circulatory death | Circulatory death | Brain death | Brain death | Euthanasia |
| Warm ischaemia time | 19 min | 24 min | 0 | 0 | 0 | 0 | 0 | 21 min |
| Cold ischaemia time | 4 hrs 6 min | 3 hrs 37 min | 5 hrs 11 min | 4 hrs 14 min | 5 hrs 4 min | 14 hrs 1 min | 3 hours 58 min | 8 hrs 13 min |
| Estimated purity (%) | 98 | 80 | 98 | 98 | 95 | 90 | 80 | 100 |
| Total culture time | 3 days | 3 days | 3 days | 3 days | 3 days | 3 days | 3 days | 3 days |
| Additional notes | N.D. | N.D. | N.D. | N.D. | N.D. | N.D. | Single-Cell RNA seq. | Single-Cell RNA seq. |

| **Islet preparation** | **17** | **18** | **19** | **20** | **21** | **22** | **23** |
| --- | --- | --- | --- | --- | --- | --- | --- |
| Donor age (years) | 65 | 54 | 72 | 64 | 71 | 69 | 37 |
| Donor sex (M/F) | M | M | M | M | F | M | F |
| Donor BMI (kg/m^2^) | 23 | 24 | 25 | 33 | 24 | 24 | 28 |
| Donor HbA_1c_ | N.D. | 35 mmol/mol (5.4%) | 40.3 mmol/mol (5.8%) | N.D. | 33 mmol/mol (5.2%) | 36 mmol/mol (5.4%) | 36 mmol/mol (5.4%) |
| Origin/source of islets | LUMC | LUMC | LUMC | LUMC | LUMC | LUMC | LUMC |
| Islet isolation centre | LUMC | LUMC | LUMC | LUMC | LUMC | LUMC | LUMC |
| Donor history of diabetes? | No | No | No | No | No | No | No |
| Donor cause of death | Circulatory death | Circulatory death | Circulatory death | Circulatory death | Brain death | Circulatory death | Brain death |
| Warm ischaemia time | 16 min | 19 min | 20 min | 17 min | 1 min | 0 | 0 |
| Cold ischaemia time | 4 hours 27 min | 24 hours 31 min | 7 hours 45 min | 6 hours 29 min | 5 hours 25 min | 5 hours 21 min | 6 hours 30 min |
| Estimated purity (%) | 98 | 97.5 | 98 | 75 | 95 | 95 | 85 |
| Total culture time | 3 days | 3 days | 3 days | 3 days | 3 days | 3 days | 3 days |
| Additional notes | Single-Cell RNA seq. | N.D. | N.D. | N.D. | N.D. | N.D. | N.D. |

| **Islet preparation** | **24** | **25** | **26** | **27** |
| --- | --- | --- | --- | --- |
| Donor age (years) | 72 | 54 | 71 | 72 |
| Donor sex (M/F) | F | M | F | F |
| Donor BMI (kg/m^2^) | 33 | 34 | 29 | 30 |
| Donor HbA_1c_ | 49 mmol/mol (6.6%) | 70 mmol/mol (8.6%) | 49 mmol/mol (6.6%) | 56 mmol/mol (7.3%) |
| Origin/source of islets | LUMC | LUMC | LUMC | LUMC |
| Islet isolation centre | LUMC | LUMC | LUMC | LUMC |
| Donor history of diabetes? | No | No | No | Yes |
| Donor cause of death | Brain death | Circulatory death | Circulatory death | Circulatory death |
| Warm ischaemia time | 0 | 0 | 0 | 15 min |
| Cold ischaemia time | 15 hours 37 min | 5 hours 17 min | 5 hours 22 min | 4 hours 59 min |
| Estimated purity (%) | 95 | 95 | 70 | 70 |
| Total culture time | 3 days | 5 days | 5 days | 5 days |
| Additional notes | HbA1c ≥ 6.5% | HbA1c ≥ 6.5% | HbA1c ≥ 6.5% | HbA1c ≥ 6.5% |

Abbreviations: LUMC, Leiden University Medical Center; M, male; min, minutes; N.D., no data

**SI appendix table 2: Donor backgrounds for human pancreatic tissue**

| Donors without history of diabetes | | | | |  | Donors with a history of type 2 diabetes | | | | | |
| --- | --- | --- | --- | --- | --- | --- | --- | --- | --- | --- | --- |
| ND donors | Age | Sex  (M/F) | BMI | Organ donation |  | T2DM donors | Age | Sex  (M/F) | BMI | Organ donation | Duration of diabetes (years) |
| 1 | 21 | M | 28 | DCDIII |  | 1 | 70 | M | 22 | DBD | 13 |
| 2 | 63 | M | 35 | DCDIII |  | 2 | 72 | F | 36 | DCD III | 12 |
| 3 | 56 | M | 25 | DCD III |  | 3 | 64 | M | 28 | DCD III | 21 |
| 4 | 57 | M | 24 | DBD |  | 4 | 60 | F | 47 | DBD | 24 |
| 5 | 61 | M | 26 | DBD |  | 5 | 50 | M | 23 | DCD III | 1 |
| 6 | 42 | F | 22 | DCD III |  | 6 | 56 | M | 22 | DCD III | 3 |
| 7 | 18 | M | 20 | DCD III |  | 7 | 67 | F | 27 | DBD | 13 |
| 8 | 68 | F | 21 | DCD III |  | 8 | 53 | M | 28 | DBD | 5 |
| 9 | 73 | F | 14 | DCD III |  | 9 | 40 | M | 34 | DBD | 1 |
| 10 | 72 | F | 25 | DBD |  | 10 | 54 | F | 19 | DCD III | 2 |
| Average | 53±20 | 6/4 | 24±5 | - |  | Average | 59±10 | 6/4 | 29±8 | - | 10±8 |

Organ donation is categorized based on the Maastricht classification.

Abbreviations: DBD, donation after brain death; DCD III, donation after controlled circulatory death; M, male; min, minutes; ND, no diabetes; T2DM, type 2 diabetes

Data are represented as mean ± SD.

**SI appendix table 3:** qPCR primers and sequences

| Gene | Primer sequence |
| --- | --- |
| *ARX* | F 5’-TGAGGCTGGACTTGACCGAGGCCCGA-3’  R 5’-GCGCCGGGTGGTGCGGAGGGAA-3’ |
| *PAX4* | F 5’-AGCAGAGGCACTGGAGAAAGAGTT-3’  R 5’-CAGCTGCATTTCCCACTTGAGCTT-3’ |
| *PCSK1* | F 5’-CTATGGAGCGAAGAGCCTGG-3’  R 5’-TCAAGTGAACCAATCTGACCCA-3’ |
| *INS* | F 5’-AAGAGGCCATCAAGCAGATCA-3’  R 5’-CAGGAGGCGCATCCACA-3’ |
| *GCG* | F 5’-CAAGGCAGCTGGCAACGT-3’  R 5’-CTGGTGAATGTGCCCTGTGA-3’ |
| *SLC16A1* | F 5’- GGTGGAGGTCCTATCAGCAGT-3’  R 5’- CAGAAAGAAGCTGCAATCAAGC-3’ |
| *NKX6.1* | F 5’- CTGGCCTGTACCCCTCATCA-3’  R 5’- CTTCCCGTCTTTGTCCAACAA-3’ |
| *OXCT1* | F 5’- GTTGGTGGTTTTGGGCTATGT-3’  R 5’- AGACCATGCGTTTTATCTGCTT-3’ |

**SI appendix materials and methods**

**Immunostaining**

Primary antibodies against C-peptide (1:400; Developmental Studies Hybridoma Bank, GN-ID4-c, GN-ID4-s), glucagon (1:2000; Abcam, ab92517), Ki67 (1:200; DAKO, M7240), NKX6.1 (1:100; Developmental Studies Hybridoma Bank, F55A12-c), MCT1 (1:200; Sigma Aldrich, HPA003324), BDH1 (1:100; Sigma Aldrich, HPA030947), OXCT1 (1:100; Sigma Aldrich, HPA012047), ACAT1 (1:200; Protein Tech, 16215-1-AP), and Hoechst (1:10000; Life Technologies, H3569) were used. Secondary antibodies Alexa Fluor 488-, 568-, and 647-goat-anti-mouse, goat-anti-rat, and goat-anti-rabbit were used to their matching primary antibody hosts.

Quantification of C-peptide^+^glucagon^-^ cells, C-peptide^-^glucagon^+^ cells, and C-peptide^+^glucagon^+^ bihormonal cells was performed by counting cells stained for C-peptide^+^ only, glucagon^+^ only and C-peptide^+^glucagon^+^, respectively. The sum of these cell types was defined as “Total cells”. A total of at least 1000 cells was counted for each condition (except in one control sample in a donor without diabetes where the total cell count was 674 due to insufficient tissue), and the percentages of each cell type were calculated. The ratio of the C-peptide^-^glucagon^+^ cells to C-peptide^+^glucagon^-^ cells was calculated.

Apoptosis was assessed by TUNEL assay (Roche). Staining was quantified as a percentage by counting at least 1000 C-peptide^+^ cells and 2000 nuclei per condition. Proliferating cells were assessed by quantifying Ki67^+^ cells in at least 200 glucagon^+^ cells and 1000 nuclei per condition.

For the quantification of NKX6.1^+^glucagon^+^ cells, at least 850 NKX6.1^+^ cells were counted per condition, and the percentage of NKX6.1^+^glucagon^+^ cells was quantified.

For the quantification of C-peptide^+^OXCT1^+^ cells in pancreas samples, at least 10 islets were examined per donor with or without type 2 diabetes, and the percentage of C-peptide^+^OXCT1^+^ cells was quantified.

Fluorescein diacetate-propidium iodide (FDA/PI) staining was used for live viability imaging. Islets of at least 80% purity were cultured in 0.5, 5, and 10 mmol/L R-BHB media for 48-hours. Aliquots of approximately 100 islets were obtained from each experimental condition and washed with PBS. FDA (0.67 µg/L) and PI (4 µg/L) were added and incubated in room temperature for 30 minutes. Samples were washed with PBS and imaged using the EVOS Cell Imaging Systems microscopy and Leica fluorescence microscopy. The percentage of FDA^+^ (live cells) to PI^+^ (dead cells) was determined using a grading system by 2 independent blinded observers.

**Type 2 diabetes single-cell transcriptomics dataset**

FASTQ files were trimmed using TrimGalore (version 0.0.6, default parameters) and mapped to the *Homo sapiens* GRCh38 reference genome (Ensembl GTF version 102) using STAR (version 2.7.7a). Multimappers were filtered out using Samtools (version 1.16.1). Reads falling in the exon region were counted with FeatureCounts (version 2.0.1) and filtered by keeping cells with a minimal number of reads below 1×10^4^ , a mitochondrial gene fraction below 0.25, and a gene count above 200. Genes that were found in fewer than three cells were removed from the downstream analysis. Single-cell RNA-seq analysis was done with the Python package Scanpy (version 1.9.3). Doublets were removed using the python package scrublet (version 0.2.3) by filtering out cells with a doublet score above 0.3. Reads were normalized to 1×10^4^ and logp-transformed. The total counts, the mitochondrial gene fraction, sjdbOverhang values used in STAR, and the dataset were regressed out. The data was batch corrected and dimensionally reduced using principal component analysis (PCA) by harmonypy (version 0.0.10), followed by computing the distance between nearest neighboring cells on the Harmony-adjusted PCA space with n_neighbors set to $\sqrt{n cells}$ rounded to the nearest integer, and n_pcs set to 50. Cells were projected in a uniform manifold approximation and projection (UMAP) and clustered using the Leiden algorithm with the resolution set to 1.25. Leiden clusters were annotated by inspecting the expression of conventional pancreatic marker genes.

Differential gene expression analysis was performed with the Wilcoxon Signed Rank Test using the rank_genes_groups function in Scanpy. We considered genes with a Benjamini-Hochberg adjusted p-value below 0.05 as differentially expressed.

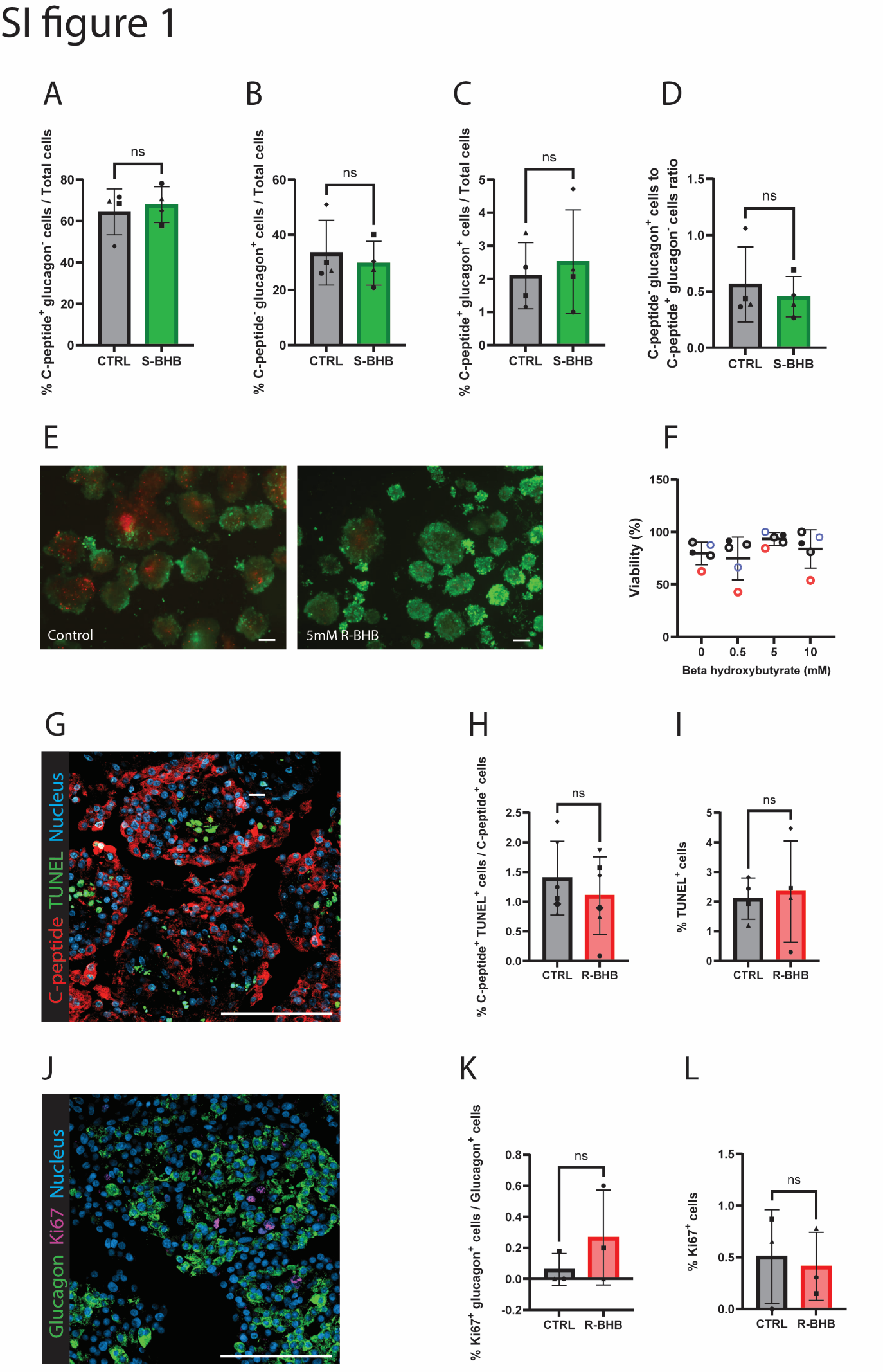

**SI appendix figure 1: Altered islet cell composition is R-BHB specific and not due to cell death or proliferation.** *A, B, C, D*: Quantification of the C-peptide^+^glucagon^-^ cell, C-peptide^-^glucagon^+^ cell, and C-peptide^+^glucagon^+^ cell proportions and C-peptide^-^glucagon^+^ cell to C-peptide^+^glucagon^-^  cell ratio of islets cultured with or without S-BHB. n = 4 donors. *E:* Representative FDA (green) and PI (red) staining of islets cultured in control and 5 mM R-BHB medium for 48-hours. *F*: Viability quantification of islets cultured in 0 (control), 0.5, 5, and 10 mM of R-BHB for 48-hours. n = 5 donors. *G*: Representative image of TUNEL (green) combined with C-peptide (red) staining. *H*: Quantification of C-peptide^+^TUNEL^+^ cells. n = 6 donors. *I:* Quantification of TUNEL^+^ cells. n = 4 donors. Data are represented as mean ± SD. *J:* Representative image of Ki67 (violet) with glucagon (green) staining. *K*: Quantification of Ki67^+^glucagon^+^ cells. *L*: Quantification of Ki67^+^ cells. n = 3 donors. Data are represented as mean ± SD. ns = non-significant. Paired Student’s t-test (*A-D, H, I, K, L*) and one-way ANOVA followed by Fishers Least Significant Difference (LSD) (*F*) was used to assess statistical significance. ns = non-significant. Scale bar 100 µm.

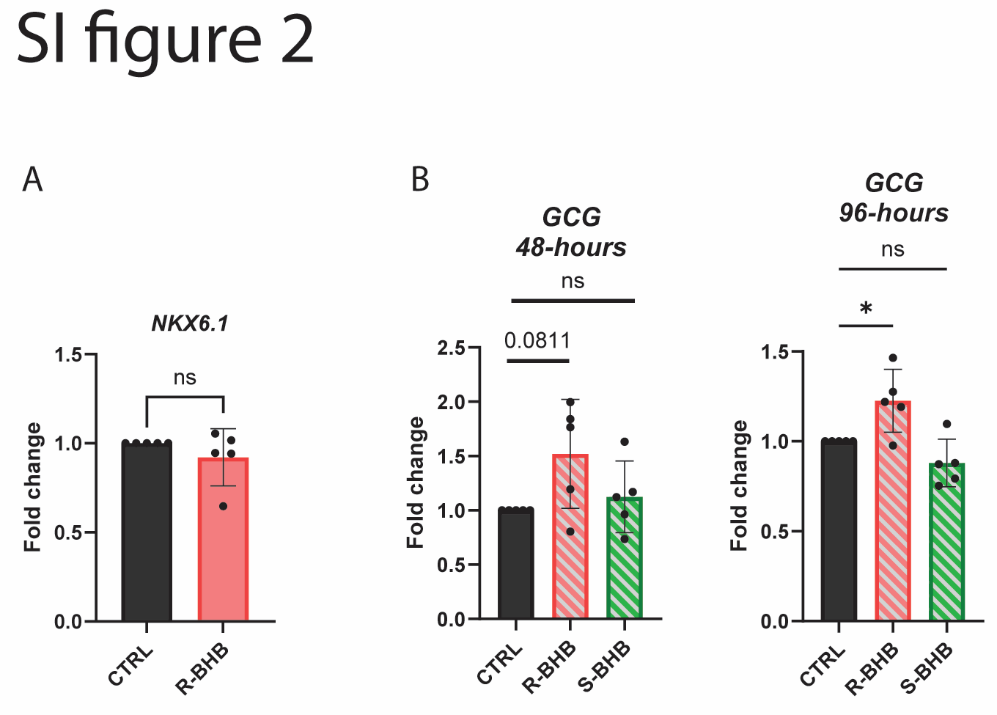

**SI appendix figure 2: R-BHB treatment does not alter *NKX6.1* expression in islets and promotes *GCG* expression in EndoC-BetaH1 cells.** *A*: qPCR analysis of *NKX6.1* expression in control and R-BHB treated islets. n = 5 donors. *B*: qPCR analysis of *GCG*  expression in EndoC-BetaH1 cells treated with R-BHB or S-BHB for 48- and 96-hours. n = 5 independent experiments. Data are represented as mean ± SD. (**p* < 0.05). Paired Student’s t-test (*A*) and one-way ANOVA followed by Fishers Least Significant Difference (LSD) (*B*) was used to assess statistical significance. ns = non-significant.

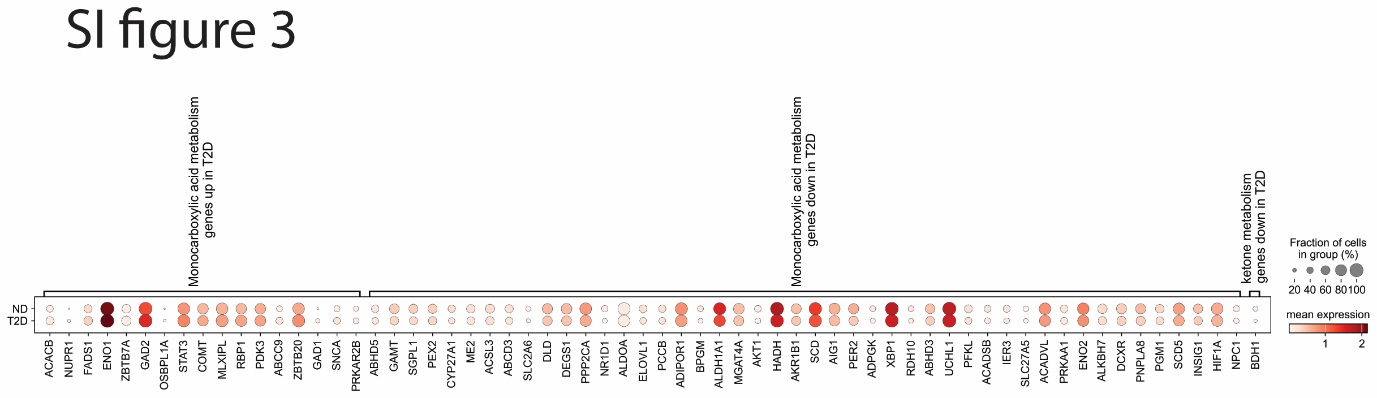

**SI appendix figure 3**

Single-cell transcriptomics analysis of significantly altered genes associated with monocarboxylic acid and ketone metabolism in beta cells from donors without a history of diabetes (ND) (n = 90) and with a history of type 2 diabetes (T2D) (n = 35).
